# How to fit in: The learning principles of cell differentiation

**DOI:** 10.1101/532747

**Authors:** Miguel Brun-Usan, Richard A. Watson

## Abstract

Cell differentiation in multicellular organisms requires cells to respond to complex combinations of extracellular cues, such as morphogen concentrations. However, most models of phenotypic plasticity assume that the response is a relatively simple function of a single environmental cue. Accordingly, a general theory describing how cells should integrate multi-dimensional signals is lacking.

In this work, we propose a novel theoretical framework for understanding the relationships between environmental cues (inputs) and phenotypic responses (outputs) underlying cell plasticity. We describe the relationship between environment and cell phenotype using logical functions, making the evolution of cell plasticity formally equivalent to a simple categorisation learning task. This abstraction allows us to apply principles derived from learning theory to understand the evolution of multi-dimensional plasticity.

Our results show that natural selection is capable of discovering adaptive forms of cell plasticity associated with arbitrarily complex logical functions. However, developmental dynamics causes simpler functions to evolve more readily than complex ones. By using conceptual tools derived from learning theory we further show that under some circumstances, the evolution of plasticity enables cells to display appropriate plastic responses to environmental conditions that they have not experienced in their evolutionary past. This is possible when the complexity of the selective environment mirrors the developmental bias favouring the acquisition of simple plasticity functions – an example of the necessary conditions for generalisation in learning systems.

These results show non-trivial functional parallelisms between learning in neural networks and the action of natural selection on environmentally sensitive gene regulatory networks. This functional parallelism offers a theoretical framework for the evolution of plastic responses that integrate information from multiple cues, a phenomenon that underpins the evolution of multicellularity and developmental robustness.

**Author summary:** In organisms composed of many cell types, the differentiation of cells relies on their ability to respond to complex extracellular cues, such as morphogen concentrations, a phenomenon known as cell plasticity. Although cell plasticity plays a crucial role in development and evolution, it is not clear how, and if, cell plasticity can enhance adaptation to a novel environment and/or facilitate robust developmental processes. We argue that available conceptual tools limit our understanding since they only describe simple relationships between the environmental cues (inputs) and the phenotypic responses (outputs) – so called ‘reaction norms’. In this work, we use a new theoretical framework based on logical functions and learning theory that allows us to characterize arbitrarily complex multidimensional reaction norms. By doing this we reveal a strong and previously unnoticed bias towards the acquisition of simple forms of cell plasticity, which increases their ability to adapt to novel environments. Results emerging from this novel approach provide new insights on the evolution of multicellularity and the inherent robustness of the process of development.

## Introduction

Organisms must sense and respond to their environment to develop, survive, and reproduce. Thus, understanding how organisms sense and respond to their surroundings has been a major subject in evolutionary biology from Darwin’s times [1]–[3]. However, during most of the 20^th^ century a simple and convenient schema in which phenotypes are environmentally insensitive (solely determined by genes) was adopted in the study of evolution [3],[4]. Accordingly, our knowledge of how the sensitivity of the phenotypes to the environment emerges from the developmental dynamics, usually known as phenotypic plasticity, is far from complete even in the simplest biologically relevant case: the living cell. Since all cells within a multicellular organism are genetically identical, phenotypic plasticity at the cell level (aka cell plasticity) is necessary for the process of cell differentiation, which in turn is crucial to build up a complex organism composed of many cell types [5],[6]. In addition, cell plasticity is also involved in the process of cell de-differentiation, a crucial event for wound healing and regeneration [7].

In cell plasticity, the presence and intensity of different environmental factors (pH, metabolites, morphogens…) determines which of a number of potential cell phenotypes will be realised [2]. From a theoretical perspective, a deterministic relationship between environmental factors (inputs) and these specific cell phenotypic states (outputs) observed in a plastic response can be described as a one-to-one mathematical function [8]. In many systems, this approach results in the so called reaction norms. While the reaction norm is a useful heuristic, it typically represents the underlying developmental processes in a simple monotonic or linear relationship between a single environmental variable and phenotype (Fig 1A) [8],[9].

**Fig 1.**
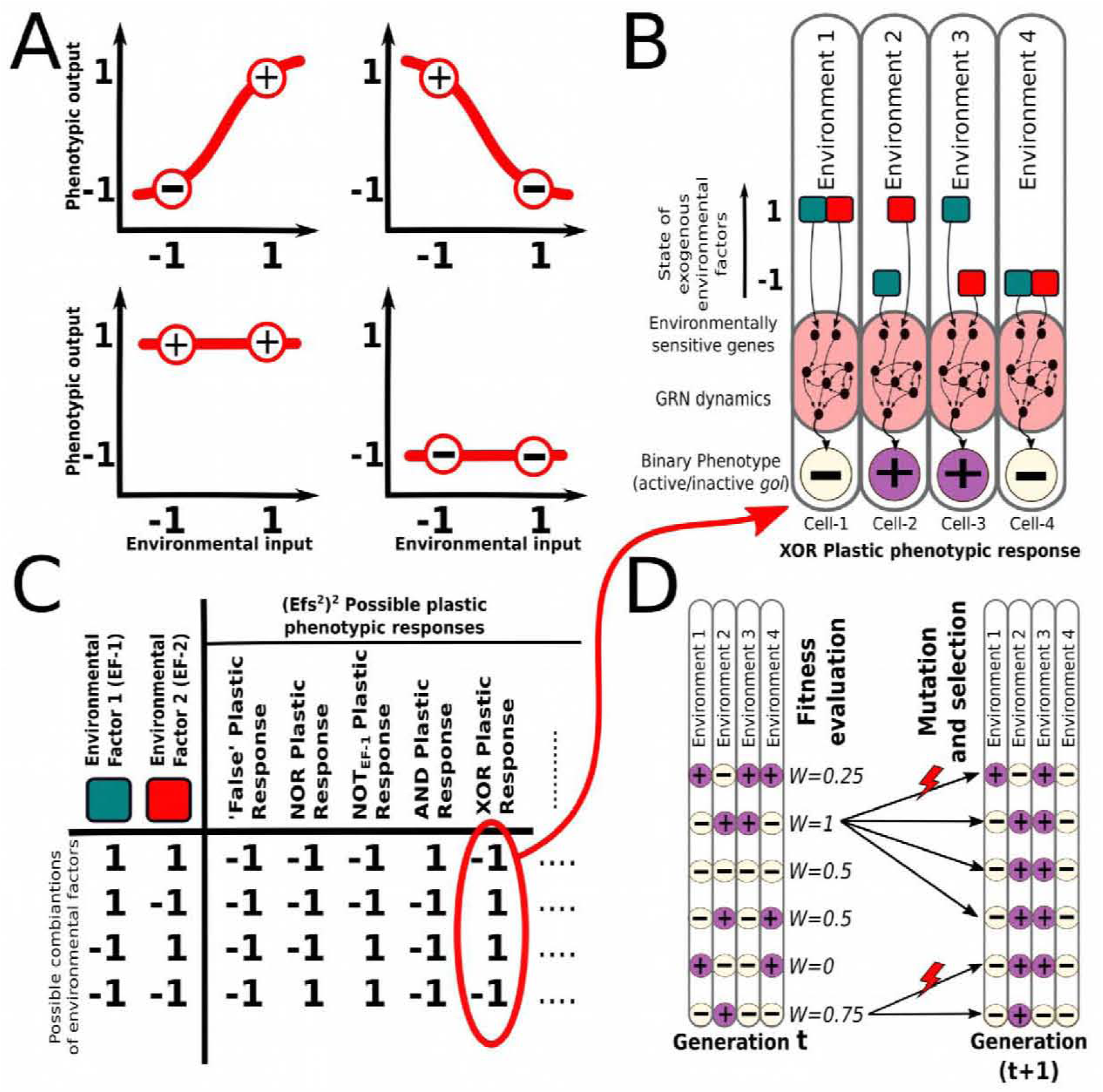
Conceptual depiction of the model. A. Models of phenotypic and cell plasticity often depict a 1-dimensional reaction norm (red line), for a single continuous environmental cue. B. In contrast, we consider different combinations of multiple discrete environmental factors (*EFs*, red and blue in the figure) determining a binary cell state. In this case, a number of possible environment-phenotype interactions can be described by means of logical functions derived from Boolean algebra (right). C. A multi-dimensional reaction norm (exemplified in the figure by the one described by the “XOR” logical function) is set as a target of a continuous GRN-based model in which the expression level of some genes is affected by the EFs. The degree of matching between the plastic response of the cells and the expected target multi-dimensional reaction norm determines the fitness (see Methods). D. Plastic cells reproduce into the next generations with a probability proportional to their fitness in a mutation-selection scenario.

In many types of phenotypic plasticity, including cell plasticity, phenotypic states are determined by more than one environmental input [1],[2],[10]. For instance, differentiating cells use a large number of signalling cascades and other mechanisms (e.g. mechano-transduction, internal biochemical clocks, etc.) as environmental inputs to gather information about when, where, and how to differentiate [7],[11]. At the cell level, the output can be described either as the expression profile of many genes or as the expression level of just a single (master) gene that controls the transcription of other downstream genes required for the cell type [10].

In order to describe the multi-dimensional (i.e., many inputs) reaction norms associated with cell plasticity, we need to use mathematical functions that generate an output (cell type), given an arbitrary number of input parameters (environmental factors like morphogens) (Fig 1C). The complete set of possible functions is familiar in the field of computer sciences and described by Boolean algebra (e.g. XOR, AND, NOR, etc.) [12]. These functions allow us to describe particular relationships between combinations of environmental cues (inputs) and the phenotypic responses (outputs): any specific multidimensional reaction norm of binary inputs and outputs is unambiguously associated with a unique logical function (Fig 1B). Furthermore, this abstraction naturally introduces a complexity measure for multidimensional reaction norms which, as we will show, offers new insights in the study of cell plasticity. Introducing logical functions is also convenient because it allows us to apply principles derived from computer science and learning theory to address some important questions regarding the evolution of phenotypic plasticity. For example, how many signals can a GRN process and how are these signals combined to generate a meaningful phenotypic response? Are all types of cell plasticity functions evolvable and are some more frequent than others? Ultimately these questions are concerned with the structure of the environment-phenotype-maps (EPMs) of cells [13]–[15]. The first aim of this work is to shed light on these questions and the nature of EPMs by characterizing cell plasticity through this novel conceptual approach based on logical functions.

In addition, our conceptual approach makes the evolution of cell plasticity formally equivalent to a simple categorisation learning process: cell differentiation requires cells to classify different combinations of environmental inputs (e.g. morphogen gradients) into a small set of possible categories (cell states). Similar categorisation experiments have been extensively studied in the context of neural networks (NN) [16]–[19]. These experiments have shown that, during the so called “training phase”, NNs exposed to a number of input-outputs can establish the logical rules that underlie the categorization process, being the rules stored in the NN circuitry [12],[20]. Then, in a subsequent “test phase”, NNs apply the same rules to novel, previously unseen inputs to generate an adequate output. This ability of any system to gather information from previous experience and to use it to offer the right response (e.g. phenotypic, behavioural, etc.) to a previously unseen challenge is the hallmark of learning [21],[22].

Whilst it seems intuitive that neural networks (NN) are capable of learning, whether the action of natural selection (i.e. random variation and selection) on gene regulatory networks (GRNs) underlying cell plasticity and cell differentiation can exhibit comparable learning capabilities is a non-trivial question. Although during the last years it has been shown that learning principles operate across a plethora of biological phenomena [21]–[26], it remains unknown if these principles apply to plastic cells performing categorisation learning tasks. Furthermore, most studies of evolutionary learning rely on simple NN-like modelling strategies, employing simplifying assumptions that are common in models of artificial neural networks but not appropriate for natural gene networks [17],[21],[26].

In this paper, our models make more realistic assumptions regarding the properties of natural gene-networks (see Methods). We also incorporate environmental inputs to these networks rather than having their inputs specified genetically, which is necessary to represent the evolution of plasticity. By doing this we demonstrate that plastic cells can store information about their evolutionary past in their GRN circuitry the same way that neural networks (NN) can store information in their network of neuronal connections [21],[22]. This feature enables plastic single cells to acquire some information about specific environment-phenotype relationships when evolving in heterogeneous environments, which are known to be commonplace in most ecosystems [20],[27]. However, in general cells might not experience *every* possible environment, so that some parts of the multi-dimensional reaction norm remain hidden – not exposed to selection [28]. We reveal that in these cases plastic cells can sometimes represent the whole reaction norm from incomplete information in the same way that learning systems can induce a complete function from partial data (generalisation) [26].

Generating an adaptive phenotypic response in an environment which have never been experienced in their evolutionary past represents a significant departure from the conventional ‘myopic’ view of natural selection, but is easily interpreted in the light of learning theory [21],[22]. We propose that such learning principles may have a pivotal role in the interpretation of cell-cell signalling of increasing complexity, in the phenotypic buffering against noisy conditions and in the evolutionary transition from single-celled free-living organisms to complex multicellular organisms [5],[29],[30]. Overall, our work suggests that the equivalence between learning neural networks and evolving gene networks holds in biologically realistic models and is thus more robust than previously demonstrated.

## Results and discussion

### Experiment 1a: Natural selection is able to find arbitrary complex forms of cell plasticity

Contrary to other approaches based on simple reaction-norms, in which all forms of plasticity are intrinsically equivalent, the use of logical functions allows us to distinguish between simple and complex forms of plasticity. This is represented by low and high Ω, respectively (see SI). Low Ω means that the phenotypic state is a simple function of one or few environmental factors (e.g. a linear one-to-one input-output map), whereas high Ω means the phenotypic state is determined by complex relationships between all inputs (e.g. a nonlinear and non-additive function of the inputs). As such, Ω is not determined by the number of environmental inputs, but by how complicated is the logic by which they are linked to the cell state [12],[31]. Each logical function is summarised in a so called *truth table*, a mathematical map which relates each combination of binary environmental factors (*EFs*) with a given binary output. Thus, we have for each function a truth table of length *2^EFs^*, which is set as a target multidimensional reaction norm.

Based on the recorded fitness over evolutionary time, our first experiment shows that cells are able to acquire multi-dimensional reaction norms of arbitrary high complexity (Fig 2). Plastic cells are able to achieve either the optimum fitness (*W*=1) or a relatively high performance (*W*>0.9). However, results also suggest that there are strong dependencies on the three parameters considered.

**Fig 2.**
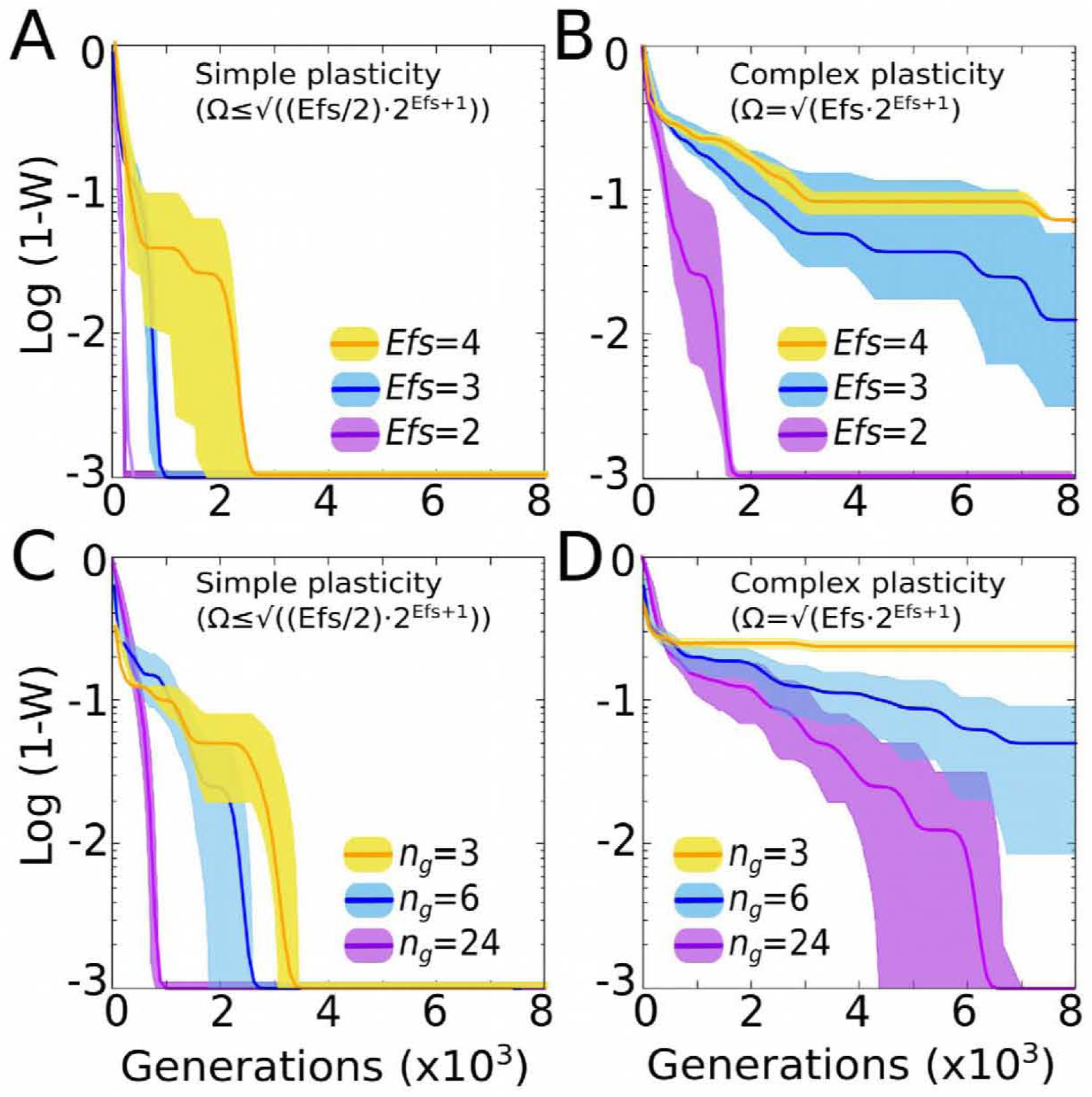
Plastic cells are able to evolve forms of plasticity of arbitrary complexity. Left panels: Different environmental inputs can be easily integrated in a phenotypic response by means of a simple forms of cell plasticity (e.g. AND, NOT functions). Evolving cells can also evolve complex forms of complexity (e.g. XOR, XNOR functions), but they take longer to do this. In general, evolving a specific form of cell plasticity is faster when the number of environmental factors involved is low (Upper panels), and/or when the size of the GRN is large (Lower panels). Below certain thresholds in the number of *EFs* and in the GRN size, (which are complexity-dependent), the system fails to accurately represent the function (see yellow lines in the right panels). Each line shows the average and standard deviation over 30 replicates. *N_g_*=12 in upper panels and *EFs*=3 in lower panels.

For instance, the amount of time required to adapt increases rapidly as the environmental dimensionality increases (See upper panels in Fig 2). Furthermore, there exists a minimal GRN size required to generate a given multi-dimensional reaction norm. This is likely because, contrary to neural networks (NNs), GRNs have strictly positive states (a gene concentration can never be negative) which prevents a straightforward digital-like computation of the inputs (see Discussion). The results also suggest that this minimal size depends on the complexity of the function required; the more complex the function the larger the network required to achieve it. All else being equal, plastic cells take longer to evolve complex multi-dimensional reaction norms than simpler ones.

This initial experiment suggests that, given enough time, biologically plausible GRNs above a certain size are capable of producing any relationship between environmental inputs and phenotypic states. In other words, with the appropriate selective pressures plastic cells are able to represent arbitrarily complex functions including those with maximal Ω=√(*EFs* ·2^EFs+1^) (Fig 2D and S1). Although we have demonstrated this capability for just four environmental factors, the observed pattern suggests that complex functions with *EFs*>4 could also be attained provided large enough evolutionary timescales (Fig 2B).

### Experiment 1b: An alternative mechanistic implementation of cell plasticity enhances evolvability

There are at least two logical ways in which an environmental signal can affect the GRN dynamics: by affecting the gene expression (either by activating or repressing it), or by modifying the strength of regulatory interactions. In this work, we have explored these two possibilities, which are referred respectively as the classical and the tensorial implementations (see Methods). Recent works suggest that biochemical mechanisms consistent with the tensorial implementation are common in nature and are involved in biological phenomena like proteins with intrinsically disordered domains [32] or non-deterministic GRNs [33]. Other examples include temperature, which causes huge phenotypic effects by affecting the morphogen diffusion or the ligand-receptor kinematics, but without altering the cellular concentration of these elements. It is worth noting that in real environments, factors that affect the strength of regulatory interactions coexist with factors that affect regulatory mechanics (e.g. pheromone-like chemicals).

In order to assess if the mechanistic manner in which the environment informs development can affect the evolution of cell plasticity, we reproduce the results of experiment 1a but by modifying the strength of regulatory interactions. As Fig 3 indicates, the results from Experiment 1a hold qualitatively true for both classic and tensorial implementations. However, cells equipped with tensorial GRNs can evolve any form of plasticity much faster than cells with traditional GRNs. Results also show that the greater differences in the adaptive rate between the classic and tensorial implementations occur for complex forms of plasticity (Fig 3B). This can be explained by considering that in complex logical functions the phenotypic effect of one environmental factor *EF1* is dependent on the state of *EF2* (higher level correlations). This dependency is naturally captured by the tensorial implementation, where the *EFs* act as modulators of the effect of other *EFs* (and genes) rather than as determinants of genetic states (Fig 3A). In addition, the dependence between the time required for adaptation (i.e., reaching a fitness of *W*=0.95) and GRN size becomes more linear under the tensorial implementation, thus relaxing the necessity of very large GRNs to generate complex form of plasticity (Fig 3B, red lines).

**Fig 3.**
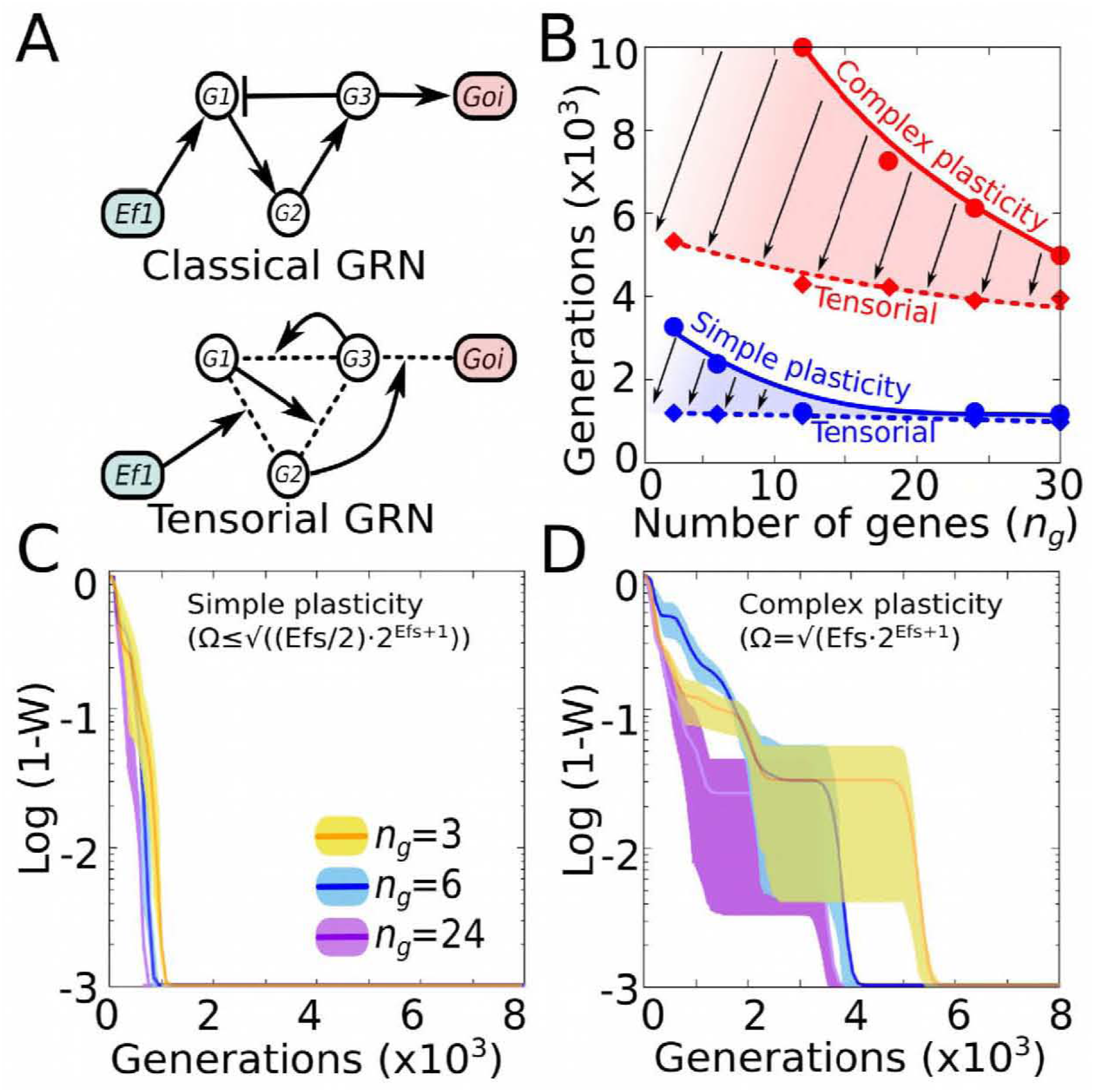
Tensorial implementation of cell plasticity enhances evolvability. A) Contrary to “classical” GRNs in which the network topology remains fixed over developmental time, Tensor-based GRNs have dynamical topology: each gene-gene interaction strength is determined by the concentration of other genes (g1, g2, etc) and environmental factors (*EFs*). In both cases the final phenotype is recorded as the binary state of a gene of interest (*goi*). B) GRNs with dynamical topologies (dashed smoothed lines) exhibit higher evolvability than classical GRNs (solid smoothed lines): lines show the average time required for reaching a fitness of *W*=0.95, which decreases for both simple and complex forms of plasticity (although the time reduction is larger in the later case, red lines and reddish shadow). C-D) As in the classical implementation, evolving a specific form of cell plasticity is faster when the size of the GRN is large, although this relationship is much more linear (compare with Fig 2C-D) *EFs*=3 and 30 replicates for all panels.

Overall, the results summarised in Fig 3 suggest that whenever the environmental factors affect the strength of gene-gene interactions, rather than the gene concentrations, the ability of biological systems to evolve complex forms of plasticity improves.

### Experiment 2: Simple forms of cell plasticity are far more abundant than complex ones

One of the possible interpretations of the results of the first experiment is that evolving cell plasticity associated with complex functions is more difficult because the number of GRNs performing these functions are scarce. That is, given any GRN generating a particular multi-dimensional reaction norm, most of its mutational neighbourhood will most likely produce either the same form of plasticity or a simpler one. In order to confirm this hypothesis, we performed an unbiased scanning of the parameter space to check how the different types of plasticity are distributed over the theoretical space of all possible GRNs and how abundant each type is (see Methods). The results, summarized in Fig 4A, show that indeed most of all the possible GRNs do not exhibit cell plasticity at all. From the subset of GRNs that show plasticity, we observe that complex plastic response functions are one order of magnitude less frequent than simple plastic response functions. Thus, the relative frequency of a given multi-dimensional reaction norm is inversely related to the complexity of the logical function which describes it. The histogram in the Fig 4A shows that the frequency-complexity relationship approximates a logarithmic function. Notice that these different frequencies do not arise from the relative frequency of each type of function in the mathematical space of all possible logical functions (their probability functions are plainly different, see dashed line in Fig 4A), but emerge as a derived property of GRN dynamics.

**Fig 4.**
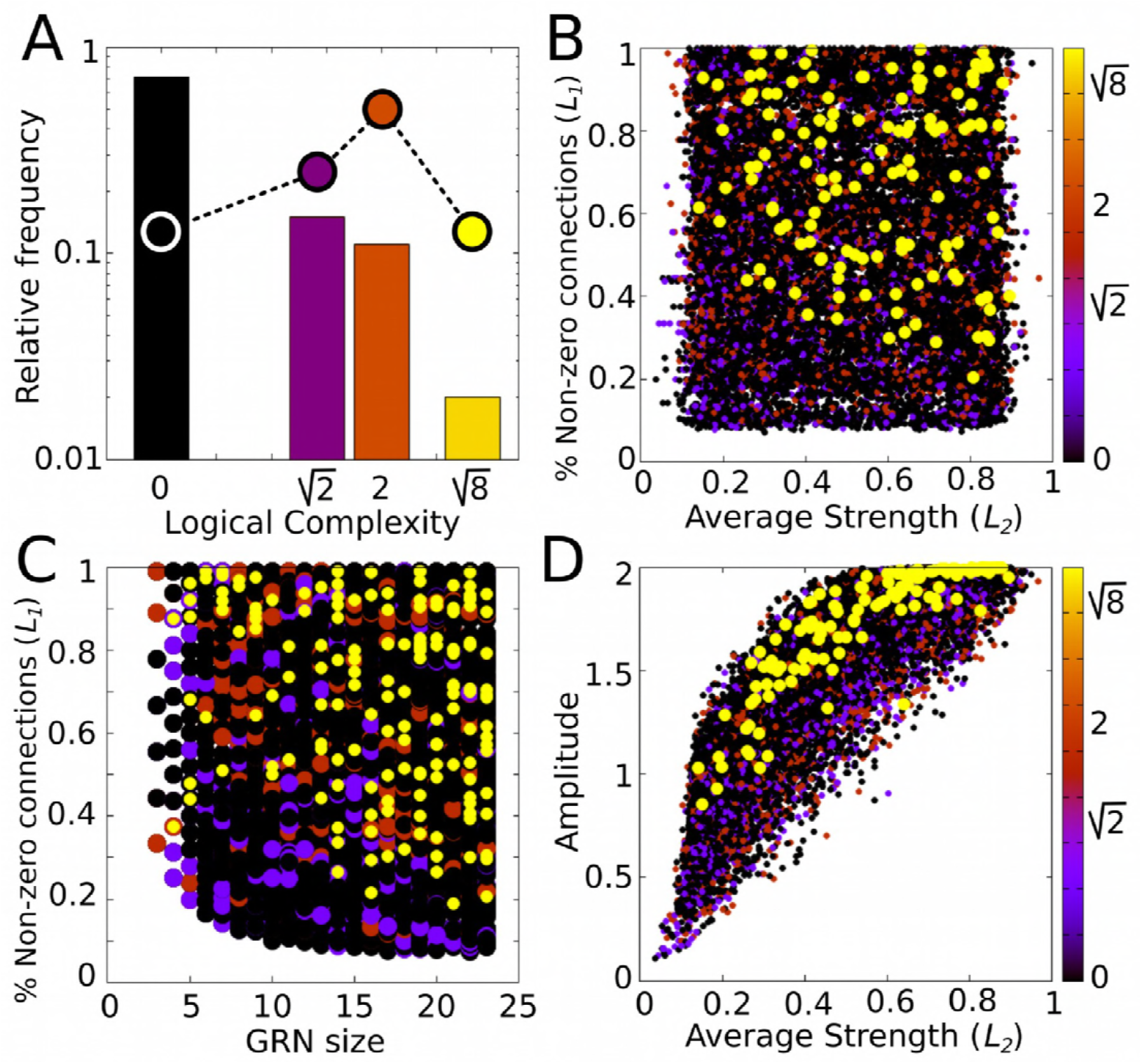
Exploring the morphospace of cell plasticity. A. (Log) Relative frequency of different types of cell plasticity according to their complexity Ω in a vast GRN space. Black column: no plasticity (the phenotypic state is purely determined by genes); purple column: phenotypic state directly determined by just one of the *EFs*; orange column: phenotypic state determined by simple combinations of the *EFs* (linearly decomposable functions) and yellow column: complex forms of cell plasticity associated with non-linearly decomposable functions (XOR, XNOR; see SI). Dashed line and dots represent the relative distribution of each family of logical functions in the mathematical space. We see that although the number of simple and complex functions that exist is approximately equal, GRNs produce simple functions much more often. B. Complex forms of plasticity (yellow dots) arise preferentially in densely connected networks with strong gene-gene interactions. CD. Complex forms of plasticity require minimum GRN sizes and disparate (0≪|*B_ij_*|<1) values in the GRN connections. In B-D panels, the % of non-zero connections and the average strengths of the GRNs correspond, respectively, to the parameters *L*_1_ and *L*_2_ used for regularisation procedures (see equations 10 and 11 in Methods). In general, less complex functions with Ω≤2 are evenly distributed in the parameter space, and are not associated with specific GRN topologies. (*EFs*=2; ≈10^5^ points).

We also examined the sub-volumes of the parameter space where each type of multidimensional reaction norms is more likely to be found. Figs. 4B-D show that, except for very complex reaction norms (Ω=√8, yellow dots in the plots), all are randomly scattered across the parameter space. The asymmetric sorting observed suggests that GRNs accounting for complex multi-dimensional reaction norms must have a minimum number of elements (Fig 4C) and dense connectivity with strong and disparate gene-gene interactions (Fig 4A and Fig 4D). In nature, selective pressures, such as a metabolic cost of keeping densely connected GRNs, may drive the system away from these regions enriched with complex forms of cell plasticity, making them evolutionarily inaccessible even if they are beneficial [26].

Together with the results from experiment 1, this bias towards the use of simple forms of plasticity seems to imply that plastic cells evolve in search spaces of reduced dimensionality: they are more likely insensitive to many of the inputs they are exposed to, and preferentially establish simple correlations between the remaining ones. This scenario of reduced search spaces has been proposed on other theoretical and experimental grounds [10],[28],[34],[35].

### Experiment 3: Simple forms of cell plasticity are evolutionary attractors

We next test whether evolutionary transitions from simple to complex reaction norms are as likely as transition in the opposite direction (isotropic, *p*(*a* → *b*)=*p*(*b* → *a*) ∀ Ω_a_>Ω_b_). The results show that, in general, evolutionary transitions are not isotropic (Fig 5A): the number of generations needed to evolve simpler reaction norms is smaller than the number of generations needed to evolve more complex reaction norms *p*(*a* → *b*)>*p*(*b* → *a*) ∀ Ω_a_>Ω_b_). Thus, although experiment 2 shows that complex forms of cell plasticity can evolve, experiment 3 shows that evolutionary transitions to simpler forms of cell plasticity are generally favoured (i.e, they can happen in a smaller number of generations, all else equal). In general, our data suggest that, given two different multi-dimensional reaction norms *a* and *b*, the speed of transitioning between them is proportional to the differences between their associate complexities (Ω_a_−Ω_b_). If both reaction norms belong to the same complexity class (Ω_a_=Ω_b_), the speed of transitioning between them is inversely proportional to their complexity class (Fig 5A, looped arrows). More formally: *p*(*a* → *b*) α Ω_a_ - Ω_b_ ∀ Ω_a_ ≠ Ω_b_ and *p*(*a* → *b*) α Ω_a_^−*k*^ ∀ Ω_a_ = Ω_b_.

**Fig 5.**
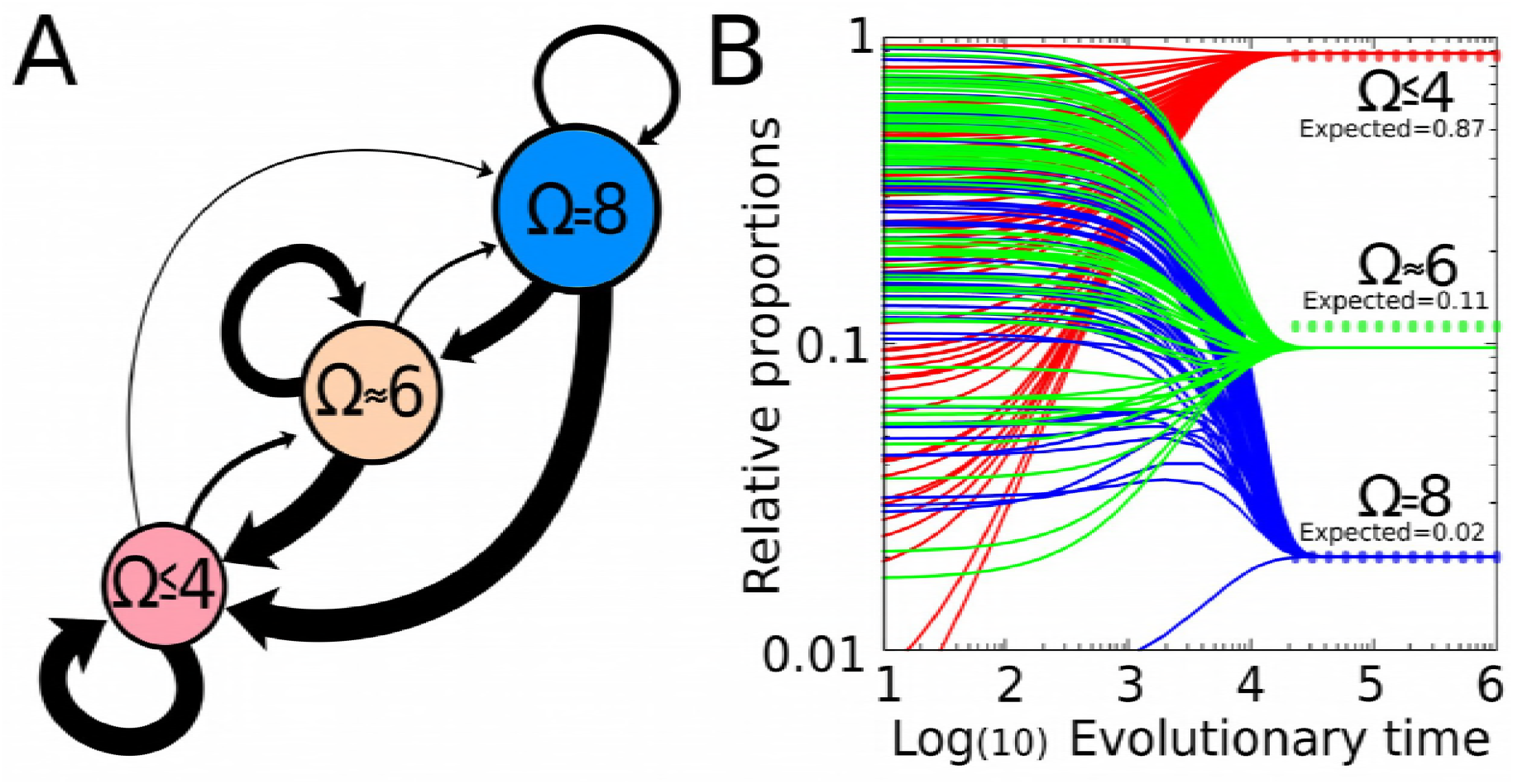
Evolvability assays. A. Transitions from complex forms of cell plasticity to simpler ones are evolutionary favoured. The width of each arrow is inversely proportional to the number of generations required for the transition from a specific multi dimensional reaction norm to another one (the wider the arrow the faster the evolutionary transition). The area of each circle is proportional to the average evolvability of each multi dimensional reaction norm (see Methods). B. Predicting the relative frequency of each type of cell plasticity from transition probabilities. Numerical simulations for the expected long-term distribution of each type of plasticity derived from the probability transitions shown in A (see equation (5) in Methods). Convergence, steady state values (0.8833, 0.0966 and 0.0201) are similar to those expected (dashed lines) according to the random scanning of the GRN space (Experiment 2), suggesting that the differential evolvability (the different probability transitions between different forms of cell plasticity) emerges from the drastic differences in the relative frequency of each type of cell plasticity in the GRN space.

The average number of generations required to go to *any* other reaction norm (including those belonging to its own complexity class) is marginally increased for complex multi-dimensional reaction norms compared to simple reaction norms (See relative sizes of the circles in Fig 5A). Thus, there appears to be a consistent evolutionary bias towards evolving simpler multi-dimensional reaction norms irrespective of the initial reaction norm considered.

We hypothesize that the differences in transition probabilities between forms of cell plasticity emerges from the relative frequency of each type of cell plasticity in the GRN space (see Experiment 2). We tested this by numerically calculating the expected long-term distribution of each type of plasticity derived from the probability transitions shown in Fig 5A. The calculations show that, when plastic cells change from one multi-dimensional reaction norm to another according to the probability of that transition (found in Experiment 3), the relative frequency of each type of cell plasticity converges to a steady-state, equilibrium value over long evolutionary timescales (Fig 5B). These values are almost identical to those yielded by the random scanning of the GRN space (Experiment 2), suggesting a causal connection between the probability transitions and the relative abundance of each type of cell plasticity (notice that these values come from different experimental setups, so they could be different).

Together, experiments 1 to 3 demonstrate a strong and previously unappreciated inductive bias towards establishing the simplest form of cell plasticity. These simple forms of plasticity are exemplified by phenotypic responses which are triggered by simple (e.g. linear, additive) combinations of the environmental cues.

### Experiment 4a: Plastic cells perform adaptive generalisation of simple plastic responses

In this experiment, we test whether cells exhibiting different classes of cell plasticity are able to use their past experience to better adapt to a new environmental challenge (that is, if they exhibit some learning capabilities [22]). We recreate classical categorisation experiments, widely used in learning theory [12],[26], by which the system has to exploit regularities observed in past situations to offer appropriate responses in novel cases (generalisation). For plastic cells, generalisation means that cells are able to infer the whole multi-dimensional reaction just by being exposed to a fraction of it (just some input-output relationships). We know from experiment 1 that plastic cells can learn any reaction norm when they are exposed to all of its points, so we now check if cells can also learn and reproduce any target reaction norm when there is missing information. In the approach we follow here, a given multidimensional reaction norm contains *2^EFs^* points, which are summarised in a truth table of the same length (see Methods and Fig 1A). To model a scenario with missing information, we set a training phase in which evolving cells can only sense a random fraction of the complete truth table. The “training set size” (TS) denotes the number of environments (rows of the truth table) that are available for cells during the training phase (*TS* < 2^*EFs*^).

As Fig 6A-C shows, the ability of plastic cells to generalize depends on the complexity of the target multi-dimensional reaction norm. When cells are exposed to a fraction of all possible environments, they interpret the environmental correlations in the simplest possible manner. That is, when some information is missing, cells evolve the simplest forms of plasticity compatible with the input-output relationships that they had experienced during the training phase. This finding is in agreement with the bias shown in our experiments 1-3 (see points with Ω<4 in the left side of Fig 6C). This inductive bias towards simplicity is advantageous when the target reaction norm has low complexity (Fig 6A): since plastic cells generalise a simple reaction norm in unseen environments, they have greater chances to fit the adaptive (target) function. Thus, under some circumstances (low complexity functions), the inductive bias exhibited by plastic cells allows them to perform better than using random responses in novel environments (Fig 6A and S1 Fig, blue line above the green line). By random responses we mean that for every point of the reaction norm for which the system has no previous information, the phenotypic response has equal probability of being +1 or −1.

**Fig 6.**
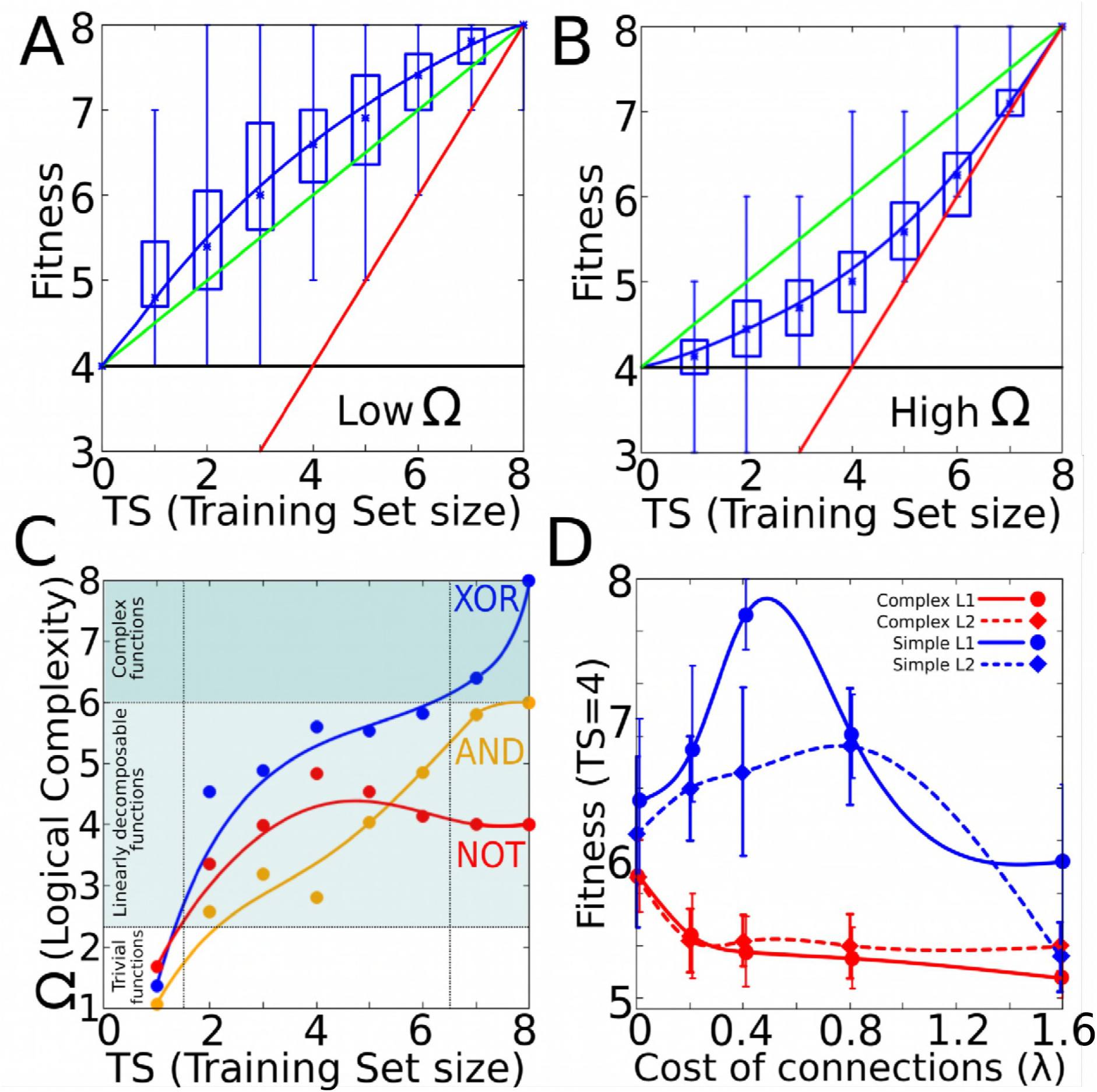
Generalisation experiments. A. When cells evolve simple forms of plasticity, they are able to generalise, performing better than chance in previously unseen environments (red line represents the information provided (*x*=*y*), green line represents the expected performance at random (2^*EFs*^−*TS*)/2; and blue line the degree of matching between the resulting phenotypes and the expected ones for a given function). Notice that blue line runs consistently higher than the green line (random response). B. Similar experiments yield poor performances when cells have to evolve complex forms of plasticity (blue line below the green one, see main text). C. When cells have incomplete information about all possible environments (left), they acquire preferentially simple forms of cell plasticity. Acquiring complex forms of cell plasticity beyond linearly-decomposable functions (see SI) requires full information (blue line). D. Generalisation experiment under *L*_1_ and *L*_2_ regularization. For this experiment, plastic cells are only exposed to half of the possible environments (*TS*=4), that is the point where their inductive bias makes the result most different from randomness. We record how the ability of cells to generalise (to approach the maximum fitness of *W*=8) changes as the cost of connections for the GRN increases (see Methods). We do this for *L*_1_ and *L*_2_ regularisation procedures (solid and dashed lines respectively) which favour sparse and weak connections respectively. We also show data for both complex (Ω>6, red lines) and simple (Ω<4, blue lines) plastic responses. The plot shows clearly how including a cost of connections improves drastically the performance of cells in generalisation experiments when cells are asked are evolved towards simple forms of plasticity. For all panels *EFs*=3; n=30 replicates, standard deviation as boxes, min and max values as error bars.

On the contrary, when the structure of the problem requires fitting complex (Ω>6) multi-dimensional reaction norms, cells cannot accurately predict the phenotypic response required in the new environments (blue line in Figs 6A-B).

In both simple and complex cases, the differences between the performed plastic response and the random response are larger for *TS*=2^*EFs*^/2, that is when the system can access just half of the available information. Obviously, also in both cases the adaptive capabilities of learning cells improve when the size of the training set increases, thus providing more information to cells, up to the limit case in which *TS*= 2^*EFs*^, which was the scenario explored in experiment 1.

Notice that for these experiments, the target plasticity function for each replicate was drawn from a subset of functions, i.e. those exhibiting the desired Ω, not from the whole ensemble of all possible logical functions of *EFs* inputs. Therefore, the tendency of cells to evolve simple reaction norms in this experiment cannot be attributed to their large statistical availability, but rather to inductive biases resulting from GRN dynamics.

In summary, when the adaptive phenotypic response is a simple function of the environmental inputs, cells will produce adaptive phenotypes in the new (previously inexperienced) environments better than expected by random completion of previously unseen rows of the truth table (Fig 6A, S1 Fig). This feature does not derive from previous selective pressures alone (i.e. it goes beyond rows that have been observed by natural selection in the past), but can be viewed as resulting from an inherent bias of evolved GRNs. That is, from the set of plasticity functions that are compatible with past selection, evolved GRNs will ‘over sample’ simple plasticity functions compared to complex ones.

### Experiment 4b: Learning improves developmental stability in a multicellular context

In complex multicellular organisms, the phenotypic state of the individual cells is determined by a complex combination of environmental cues and endogenous signals such as morphogen gradients [7]. In order to produce non-trivial and spatially organized developmental patterns, differentiating cells need to respond appropriately to these complex signals. In principle, it could be the case that every possible response (cell-state) has been the explicit target of past natural selection. However, the previous experiment suggests that this is not always necessary. If the target pattern results from simple morphogen combinations and natural selection has occurred over enough morphogen-phenotype combinations, the evolved GRNs will fill-in remaining regions of the multi-dimensional reaction norm better than by chance. To illustrate this generalisation capacity in a multicellular differentiation context, we set a 2D embryonic field of 1250 cells which can gain positional information from the asymmetric spatial distribution of a few different morphogens (Fig 7). The morphogens are spatially distributed along simple gradients, but none of them alone contains enough information to produce the final developmental pattern. The pattern emerges by integrating the morphogenetic cues according to a relatively simple composite logical function (Fig 7A).

**Fig 7.**
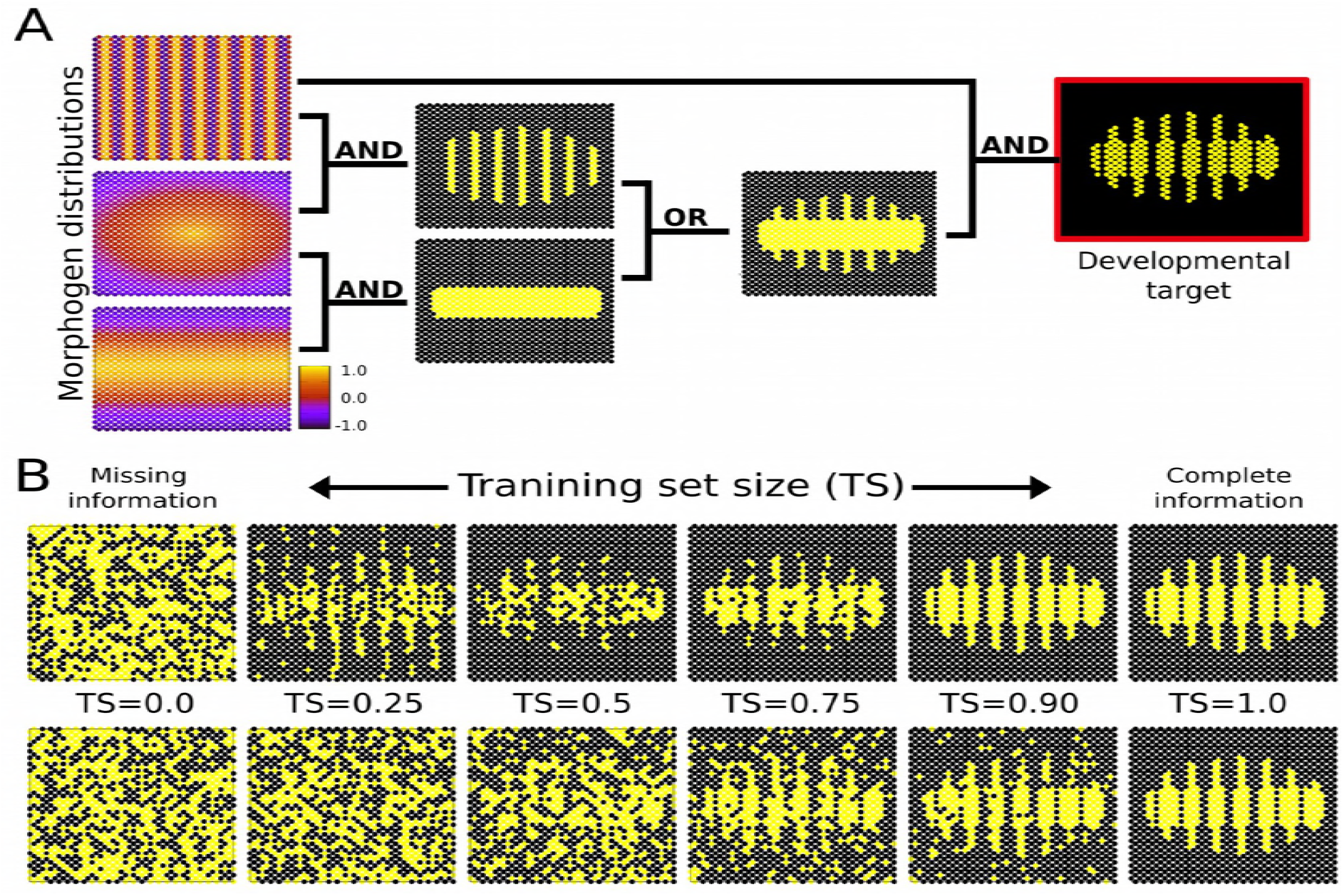
Developmental implications of learning-based cell plasticity. A) Simple spatial distribution of three morphogens over a two-dimensional (50x25) field of cells (similar gradients are commonly found in the early developmental stages of many organisms). In this example, these three environmental signals are integrated by individual cells (according to relatively simple logical functions) to form a segmented developmental pattern. The complete plasticity function is represented by a truth table of 32 rows specifying whether the relevant cellular output should be on or off for each possible combination of morphogens (see Methods). B) Individual cells are evolved under selective conditions that expose them to some rows of this table but not all (see Methods). Under this scenario of missing selective information, cells equipped with real GRNs perform by default a simple logical integration of the morphogen cues, which in this example results in a much more robust and uncertainty-proof developmental process (upper row). When the training set contains all the information necessary to completely define the plasticity function, natural selection finds a GRN that calculates this function accurately (right). Of course, when past selection contains no useful information, the phenotype of the GRN is random (left). In between we can see that the generalisation capability of the evolved GRN ‘gives back more than selection puts in’-i.e. the phenotype produced (top row middle) is visibly more accurate than the training data experienced in past selection (bottom row middle). This is quantified in Fig S1. In the bottom row, cells acquire any random function (not just the simpler ones) compatible with the rows experienced during past selection, which prevents generalisation (see main text).

When individual cells are evolved to perform such a function (complete information) during the training phase, they accurately recreate the complete segmented pattern during the test phase. However, it can be the case that individual cells evolve in selective conditions that are missing information – i.e. where just a random subset of the multidimensional reaction norm is selected for and the remainder has not been subject to selection (see Methods). In a developmental context that means that cells receive just a subset of the morphogens they have evolved to recognise (different cells receive different “bits” of incomplete positional information). In our experiment, since the logical structure of the multi-dimensional reaction norm that generates the segmented pattern is simple, plastic cells can fit it from just a few points. Essentially, multi-dimensional reactions norms governed by simple logical functions belong to the category of problems that cells are well suited to solve because of their inductive bias towards simplicity (S1 Fig). Thus, as Fig 7B shows, recognisable patterns emerge even in very disturbed conditions. In the same figure (lower row) we can also see how plastic cells that lack learning-based buffering fill-in missing information at random. In this case, cells can acquire *any* of the remaining logical functions compatible with the experienced ones, not just the simpler ones (which happen to be the adaptive ones in this example). This experiment shows that in the absence of any inductive bias, plastic cells are generally unable to repair corrupted phenotypes (Fig 7B, S1 Fig).

Comparable noisy conditions are known to be commonplace in nature, and biological systems have developed a variety of mechanisms to cope with them [36]. Our experiment suggests that under these non-ideal conditions, learning may act as one of these buffering mechanisms. However, unlike other phenomena that increase developmental robustness by reducing the overall sensitivity of the system to the external conditions, learning can increase developmental robustness while keeping the environmental sensitivity fully functional. This apparent paradox is explained because the buffering process described here is concerned with *correlations* between morphogens rather than by the number or absolute concentrations of morphogens. In other words, this mechanism does not have to reduce the number of morphogens or the sensitivity of the system to these morphogens, it works by restricting the number of ways in which these morphogens can be combined to generate phenotypic outputs.

### Experiment 5: Costly connections amplify the performance in generalisation experiments

In this experiment we explore some conditions which may enhance generalised phenotypic responses. In the light of Experiment 2, there seems to be a dependency between the complexity of the plastic phenotypic responses and the GRN topology (e.g., complex multi-dimensional reaction norms are produced more often by plastic cells having densely connected GRNs). Thus, we examine an evolutionary scenario in which the generalisation experiments must be performed by GRNs with a particular topology. Specifically, we force GRNs to acquire a certain topology (e.g. a weak connectivity) while evolving in the training phase of a generalisation experiment. We do this by implementing a connection cost in the evolving GRNs (see Methods), so that the connectivity of the GRN contributes to the cell’s fitness. Introducing such a cost is biologically meaningful since densely inter-wired GRNs are known to be selected against for a number of reasons, including the intrinsic metabolic cost of synthesizing a larger number of organic molecules, the lack of efficiency in the substrate-ligand interactions between gene products derived from thermodynamic considerations, and because the GRN dynamics is more likely to exhibit chaotic behaviour [26],[37]. Thus, although there exist some evolutionary advantages in having redundant elements in the GRN (e.g. increased robustness, [15],[36], the selection against too complex GRNs is biologically consistent. In addition, introducing a connection cost (aka parsimony pressure or regularisation) is a widely used procedure in learning theory and artificial intelligence that is known to alleviate the problem of over-fitting [12],[21],[26], thus providing us an opportunity of testing the formal correspondence between learning principles and cell plasticity.

We implemented two different types of costly connections, one (*L*_1_) that favours sparse connectivity in the GRN and one (*L*_2_) that favours GRNs with weak connections. As Fig 6D shows, both procedures have a qualitatively similar effect on the ability to generalise. However, the effect depends strongly on the complexity of the multi-dimensional reaction norm that cells have to evolve. For complex forms of plasticity (red lines in Fig 6D), introducing a cost of connections makes the generalisation response worse (approaching the minimum value of TS=2*^EFs^/2*=4, where the system output contains just the information provided during the training stage). This happens because introducing a cost in the GRN connections pushes the system towards regions of the GRN space enriched with weakly and sparsely connected GRNs (where no complex forms of plasticity can arise), so the system cannot fit the complex target (known as “underfitting” [26]).

The opposite is observed when the plastic cells generalise over simple forms of plasticity. In that case (blue lines in Fig 6D), having sparse and weakly connected GRNs increases the likelihood of cells to perform simple forms of plasticity, which increases their performance under previously unseen environmental conditions (i.e., generalisation). However, when the connection cost becomes too high (beyond an optimum λ of 0.5 for *L*_1_ and 0.8 for *L*_2_), the enhanced performance disappears. That is probably related to the fact that even to perform the most basic computations, GRN require a minimally dense topology: extremely simplified GRNs do not perform any kind of computation at all.

## Conclusion

In this work we have introduced a novel theoretical approach for the study of cell plasticity that exploits a formal mapping between cell plasticity and logical functions. Under this approach, the different multi-dimensional reaction norms exhibited by plastic cells are associated with specific logical (mathematical) functions derived from Boolean algebra. This idealisation enables us to measure the complexity of the different forms of cell plasticity.

By using this approach, we have shown that plastic cells are able to display arbitrary complex forms of plasticity such as those associated by hard-to-compute, non-linearly decomposable logical functions (Ω=√((*EFs/2*)·2^EFs+1))^. This class of logical functions is well known in learning theory because they cannot be learned by linear models (such as the linear perceptron [12]). To learn such functions requires the ability to represent and learn correlations between the inputs or to compare one input with another (e.g. inputs have different values, XOR), a feature which is mandatory for associative learning [12],[21]. In computer science the ability of artificial neural networks to represent and learn such functions is well understood. In contrast, whether biologically realistic GRNs with an unstructured topology (e.g. without explicit hidden layers) and strictly positive variables ([*g_i_*]>0) can be evolved to represent such functions has not been previously shown. The implementation of learning theory approaches to study phenotypic evolution has been developed in prior work [21],[22],[24],[26]. That work, however, is concerned with the evolution of evolvability (generating phenotypic variation, that may be adaptive, given new genetic mutations) rather than plasticity (adaptive phenotypic responses to environmental cues). This paper thus focuses on how evolving systems rely on learning principles to classify a series of elements (e.g. combinations of morphogens) into a set of discrete categories (e.g. cell state).In learning terms, the former corresponds to a generative model whereas the latter, addressed in this paper, corresponds to a discriminative model [12]. Our approach represents therefore a novel conceptual tool in the study of cell plasticity.

Experimental evidence from the field of synthetic biology shows that *in vivo* circuits (e.g. DNA-based logical gates or genetically engineered metabolic networks) are able to solve simple problems such as non-correlational AND-like functions (Ω=√(*EFs*·2^EFs+1^)). For more complex functions, the complexity of the biological circuits rapidly escalates with the problem complexity in a way that makes empirical approaches very challenging [38],[39]. In those studies, the authors suggest that the lack of scalability is caused by the fact that real GRNs have to reuse the same genetic modules. In the present work, we have demonstrated that complex problems can be solved by biological systems if they are endowed with larger, denser and more strongly connected circuits (Fig 2). A network with signed states (e.g. [21]) can compute the same function using fewer nodes than a network with unsigned states (i.e., [*g_i_*]>0). This may explain why real GRNs, which can only have non-negative gene concentrations, must be large and densely connected in order to perform non-trivial computations [12]. This finding could be useful to enable engineered cells in synthetic biology to address more complex problems.

Other computational models of plasticity avoid the complications created by nonnegative states by following different strategies. Some introduce abstract state variables representing ‘gene expression potentials’, which are signed, rather than gene-expression levels, which are non-negative [21],[26]. Others make use of an output layer to convert unsigned gene-expression states into signed phenotypic traits [40]. The most common approach is to directly encode the reaction norm parameters (e.g. slope) in genetically heritable variables, thus bypassing the generative dynamical processes responsible of the inductive biases [8]. Our results show that a conventional GRN model with unsigned states is sufficient to evolve arbitraily complex forms of adaptive plasticity.

Specifically, we demonstrated the potential of GRNs to generate complex multidimensional reaction norms under two biologically plausible scenarios: one in which the environmentally-sensitive factors are gene expression levels (classical GRNs) and one in which the environmentally-sensitive factors are the strengths of gene-gene interactions (tensor-based GRN). We show that the tensor-based implementation greatly improves the capacity of cells to evolve adaptive forms of plasticity, especially the more complex ones (i.e., those highly non-linear input-output maps). This observation suggests that more complex forms of cell plasticity can be expected in response to particular factors, such as temperature, that influence expression dynamics through modulation of gene-gene interactions rather than individual gene-expression levels.

In addition, our results reveal that complex forms of plasticity are difficult to evolve because the hypervolumes of the GRN space containing them are very small and do not form a connected region (Fig 5B). The networks capable of complex plasticity are accordingly rare and require a dense topology of strong and heterogeneous connections. Likewise, the vast majority of new mutations in a given GRN drive the evolving cells into regions of equal or less complex plasticity. This applies to mutations that cause changes in GRN topology as well as mutations that cause changes in gene-gene interaction strengths [14]. By means of numerical simulations, we quantitatively predict the relative frequency of each type of plasticity from the probability of transitions between them (Fig 5B). This suggests that low evolvability of complex reaction norms is a consequence of the scarcity in the GRN space of the networks that generate complex reaction norms. A similar bias towards simple input-output maps has been recently proposed for artificial neural networks [19], and algorithmic theory suggests that it might be a general property of every computable functions [31]. Our contribution is to demonstrate that this bias is intrinsic to cell plasticity, and that it emerges from reasonable model assumptions of gene-expression mechanisms. This may help to explain why many forms of plasticity tend to rely heavily on a low number of environmental cues even when many cues potentially are informative.

This bias towards simple forms of plasticity is consistent with studies on genotype-phenotype-maps (GPMs) which show that more complex phenotypes are more scattered in the parameter space, thus being far less frequent than simple ones [13],[41],[42]. These qualitatively similar properties of GPMs and environment-phenotype-maps (EPMs) are likely due to the fact that they arise from the same underlying dynamical system: the GRN [14],[35].

Although further work is required in order to systematically explore how complex GPMs and EPMs emerge spontaneously from GRN dynamics (and more complex multi-level developmental systems), our work provides a theoretical foundation for what to expect when plastic cells are exposed to novel multi-factorial environments. Knowing that cells preferentially acquire phenotypic states determined by the simplest combinations of the external factors may be useful for cell cultures exposed to new multi-nutrient growth media or for cancer research where tumor cells are treated with new multi-drug cocktails [16].

In the context of cell plasticity, learning principles are relevant in three complementary ways. One is simply that by preferentially using simple logical functions, plastic (differentiating) cells do not equally consider all the available inputs (signalling molecules). Rather, cells ignore most inputs so that their phenotype depends on simple combinations of the remaining ones. This effectively decreases the size of the search space, making plastic cells evolve within a signalling environment that has an actual dimensionality much lower than the apparent one. In learning terminology, this is an inductive bias. Such biases favouring simple functions increase the chances of evolving an optimal combination of signals in a finite amount of time if the relationship between environmental cues and optimal phenotypes is, in fact, simple. For example, in the case of free-living single cells, if this bias is useful, it may enable cells to better track seasonal changes by using just a fraction of the information provided by the environment [27].

The second reason why learning principles are relevant is more subtle. Under the scenario we propose here, the differentiating cells preferentially establish simple logical relationships between the environmental factors they have been exposed to (a form of inductive bias). This inductive bias towards simple functions means that plastic cells have an intrinsic, GRN-based mechanism for solving simple problems (although we have demonstrated that this bias is not limiting given the appropriate selective pressures). Thus, whenever the unforeseen challenge is a simple problem, the intrinsic bias towards simple plastic responses enables cells to show an improved performance compared to a random response (Fig 6A, Fig 7B, S1 Fig). This bias can be even more pronounced if the GRN connections are costly, a feature which is known to be common in real cells (Fig 6D). The difference to standard notions of Darwinian evolution is not trivial. In classical Darwinian evolution, plastic cells placed in a novel environment have no information to guess which will be the right (fitter) phenotype in that context [43]. In the absence of any inductive bias which causes cells to prefer one type of plasticity function over another, the cells will blindly proceed by trial and error until the right cell state is eventually found, which may hinder or delay adaptation over evolutionary time [23].

The third reason why learning theory is relevant applies within a multicellular context: once plastic cells have evolved the cell differentiation function (which cell state corresponds to each combination of morphogens), learning can buffer the process of development against noisy or corrupted signalling pathways (Fig 7B). Plastic cells achieve this by establishing only basic correlations between a few morphogens, which makes them consistently differentiate in the same phenotypic state (cell type) even when new signalling pathways arise by mutation. This inductive bias of the GRNs may increase phenotypic robustness against environmental or genetic perturbations, along with other more known properties of biological GRNs like redundancy or modularity [7],[40]. Importantly, the learning-based robustness that we report here does not require a lack of sensitivity to the environmental inputs, but can arise from a logical simplification of the possible input-output relationships.

From a macroevolutionary perspective, it must be noted that the inductive bias towards simple forms of plasticity is present without being specifically selected for [44]. Rather, this bias towards simple plasticity is an inherent property of GRN dynamics which was already present in single-cell organisms well before the evolutionary transition towards multicellularity [5],[6],[44]. During this transition, which has happened at least 40 times [29], [30], the functional integration of the new level of complexity crucially depends on the ability of the lower-level entities to correctly interpret and respond to a vastly rich set of intercellular signals [29]. Coming from free-living states, the multi-cellular state might have been very challenging [7]. However, as we have demonstrated here, the ability of cells to generalise from simple forms of plasticity may have facilitated the ability to adapt to more complex signalling environments. Testing how inherent inductive biases caused by cellular properties may have interacted with other factors in the self-organization of differentiated cell aggregates constitutes an exciting prospect for future research [45].

A possible criticism of our study is that plastic cells can only take advantage of adaptive generalisation if the plasticity function is simple (Fig 6A, S1 Fig). However, this does not undermine the generality of this phenomenon for two reasons. First, biologically, the signalling context inherent to multicellularity was not a pre-existing ecological niche to which cells adapted, but rather an environment that cells Co-created themselves during the evolutionary transition [29],[30]. From our findings, which show that single cells exhibit a bias favouring simple plasticity functions, it seems plausible that such simplicity pervaded the signalling context of the newly created multicellular environment. In other words, cells have evolved developmental patterning which requires only simple combinations of morphogens because this matches the simple plasticity they can exhibit. It is within this constructed space of limited dimensionality where plastic cells can take advantage of the learning principles; that is, cells have created a problem that they were already able to solve. Second, logically, a bias that favours simple solutions fits well with generic properties of the natural world. This “universal logic”, which results from probability theory and is often referred as the Occam’s razor, states that all else being equal the simplest solution tends to be the correct one. The opposite bias (i.e., one favouring complex or less parsimonious solutions) would not produce effective learning in most cases.

Another criticism may be that real cells might have evolved mechanisms to overcome this bias towards simple forms of plasticity, for instance, by modifying the GRN connectivity. However, this would require an adaptive pressure towards denser GRNs, which *ceteris paribus* seems unlikely given metabolic costs of regulation [26]. A more plausible scenario, given GRN connections that are costly, results in an even more pronounced bias towards simple forms of plasticity (Fig 6D), so that this bias seems to be unavoidable.

Ultimately, the ability of plastic cells to mimic logical functions and to perform proper learning with generalisation is likely to arise from the structural, dynamical, and functional isomorphisms between GRNs and NNs: they both partition the space of possible inputs into a series of states or attractors [13],[37],[42], both are adaptable via incremental improvement principles [15],[41],[46] and both exhibit similar inductive biases [14],[31]. In addition, both exhibit increased performance under similar adaptive pressures [26]. For example, we have shown that some procedures commonly used to improve the performance of artificial NNs (such as the *L*_1_ and *L*_2_ regularisation) can also enhance the ability of plastic cells to acquire specific reaction norms from incomplete information (Fig 6D).

All these commonalities between GRNs and NNs allow us to transfer knowledge derived from learning theory to illuminate the domain of cell plasticity. This approach is ultimately possible because, put in simple terms, gene networks evolve by similar principles to those by which cognitive systems learn.

## Methods

### The core model

The developmental model is a GRN-based implementation with non-symmetric continuous (non-Boolean) unsigned state variables and explicit environmental factors. Basically, the model consists of *N_g_* genes or transcription factors that regulates expression of other genes. The state of the system in a given time is determined by the *G* vector, containing all the concentration profiles of all the *N_g_* genes and gene products. Genes and gene products have continuous concentrations *g_i_*≥0, and interact with other genes or gene products by binding to cis-regulatory sequences on gene promoters, conforming a gene regulatory network (GRN). The regulatory interactions of this GRN are encoded in a *B* matrix of size *N_g_* x *N_g_*. All genes and gene products are degraded with a decay term *μ*=0.1.

Environmental factors (*EFs*) are implemented as exogenous cues affecting the levels of gene expression by activating or repressing them. Each *i* of the *EFs* environmental factors (*EFs*<=*N_g_*) affect a single gene with an intensity of *Ei*=−1 or *Ei*=+1 (see Equation 2 below). Environmental factors are not degraded over developmental of evolutionary time (Fig 1). The GRN generates a suitable phenotype by integrating information about both the genetic state and the environment by an iterative process over a number of *Devt* developmental time steps. The single-trait phenotype is recorded as the gene expression of a gene of interest (*goi*), which is different from the environmentally sensitive genes affected by the *EFs*.

Although the model is highly simplified and it does not explicitly include space, it captures the most typical mechanisms by which real genes regulate one another. Similar models have yielded valuable insights on the evolution of plasticity [40],[47], the evolution of modularity [48], developmental memory and associative learning [21], the emergence of rugged adaptive landscapes [37], the evolution of biological hierarchy [49], or the role of noise in cell differentiation [36] (Fig 1).

The gene-gene interactions within the GRN follow a non-linear, saturating Michaelis-Menten dynamics, which is a special type of Hill function widely used for modelling biochemical binding processes [50]. Other classes of non-linear, monotonically increasing functions have been explored in previous works giving consistent results [51]. In the *B* matrix, each interaction strength *B_ik_* represent the effect of gene *j* in the transcription of gene *i*, acting either as a repressor (when −1≤*B_ij_*<0) or as an enhancer (when 0<*B_ij_*≤1). Developmental dynamics is attained by changes in gene concentration over developmental time. Thus, the concentration of the gene *i* in the cell (*g_i_*) changes over time obeying the following differential equation:

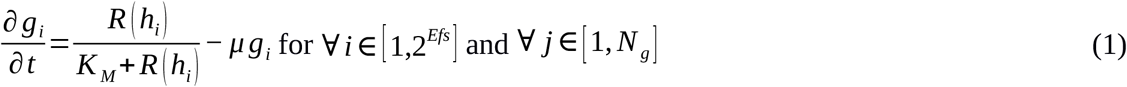

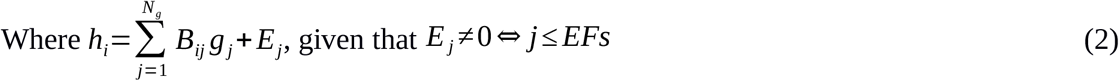

In the first term of the equation (1), R (*h_i_*) is the Ramp function (R(x)=x, ∀ x>=0 and 0 otherwise), which prevents eventually negative concentrations in gene products resulting from inhibiting interactions, and *K_M_* is the Michaelis-Menten coefficient. Without loss of generality, we set *K_M_*=1 (other choices are known not to affect the results, [41],[51]). The second term of the equation 1 describes the degradation process that affects every gene in the system. Equation 1 was numerically integrated by means of the Euler method (δ_t_=10^−2^) over a maximum developmental time of *devt*=10^6^ iterations (arbitrary time units) or until an steady state is reached. This happens when all the normalized gene concentrations remain the same within a threshold of 0.01 over an interval of 1000 developmental time units (i.e., when |G_devt_/max(*g*(*i*)_devt_)}={G_devt+1000_/max(*g*(*i*)_devt+1000_)}|≤0.01). Manually directed simulations showed that when the system does not reach a steady state within this maximum developmental time, it is because it is undergoing either cyclic or chaotic behaviour, such that it never will saturate.

Although the equation 1 has no explicit noisy term, some noise is introduced in the initial conditions in order to break the initial symmetry of the system. Specifically, in *devt*=0, all gene concentrations are set to *g_i_*=0.1+ξ, being ξ a small number randomly drawn from a uniform distribution ξ~U(−10^−2^,10^−2^). Previous works suggest that small modifications in the equations (e.g. the inclusion of a noisy term, the choice of a specific Hill-function, etc.) do not substantially alter the dynamics of these models, thus implying that the developmental dynamics actually relies in the GRN topology itself [41],[51].

In an extended version of our basic model, environmental factors do not affect particular genes, but the strength by which genes interact between them (Fig 3A). We implement this by using higher-order matrices called tensors, so this version of the model will be referred hereafter as the tensorial implementation. In here, the GRN topology itself is not fixed but dynamic over developmental time, and consequently the *h_i_* term of the Equation (2) is now described as:

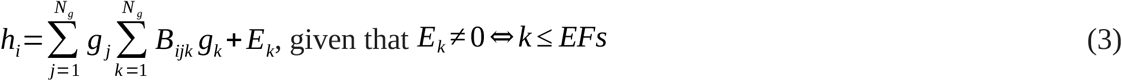

In the tensorial version, the *B_ijk_* matrix has an extra dimension “*k*”, which determines which genes contribute to the establishment of the interaction strength between the gene products *i* and *j*. In this tensorial implementation, each environmental factors (*EFs*<=*N_g_*) contributes to determine the interaction strength between different gene products with an intensity of *E_k_*=−1 or *E_k_*=+1. All the remaining elements in the tensorial version are implemented the same way as in the basic model. The phenotype characterisation and the different experimental setups described in the ensuing paragraphs apply to both the basic and tensorial implementation (Fig 3A).

In many biological examples, specific gene expression levels are known to trigger biochemical, developmental or physiological responses in a binary manner. Above a certain threshold in gene concentration (when the gene is active), the response is triggered, and below the threshold (inactive gene), it is not. Motivated by this fact, the phenotype of a cell in our model is conceptualized as the binary expression profile of the gene of interest (*goi*): *g*’_*goi*_=1 ↔ *g_goi_* ≥*δ_t_*=10^−2^ and *g*’_*goi*_=−1 ↔ *g_goi_* ≤0.1. This allows us to treat the phenotypic outcome as a boolean output.

### Experiment 1. Representations

In our first experiment, we simply evolve different GRNs towards different multidimensional reaction norms (that is, to match specific input-output maps or truth tables), including those described by complex logical functions (Fig 1) [47]. Whilst some authors have implemented this as a single population which faces sequentially all possible environments over a number of generations (each environment being a specific input-output row of the whole truth table [15],[25],[26], we split in each generation the entire population of plastic cells into a set of minimal (single-cell) isogenic sub-populations, with each subpopulation being exposed to a specific environment (i.e. a specific combination of the environmental factors). This procedure is formally equivalent to the first one but allow us to save computational resources [12] and to ensure that all environments equally contribute to the acquisitions of specific forms of cell plasticity.

Since an environmental factor can only take binary values (−1,+1), there are *2^EFs^* different environments resulting from a combination of the *EF* environmental factors. Thus, the multi-dimensional reaction norm linking the (binary) environmental inputs with the (binary) phenotypic outputs can be represented as a truth table of length 2^*EFs*^. The phenotypic output of the truth table is set as the target function *P* (Fig 1),

In each generation, the target function *P* is compared to the resulting phenotype in each environment. The sum of all the matches between the phenotype and the target *P*, scaled by the size of phenotypic vector, determines the fitness *W* of a given GRN. The fitness *W* determines the likelihood of a GRN to contribute to the next generation:

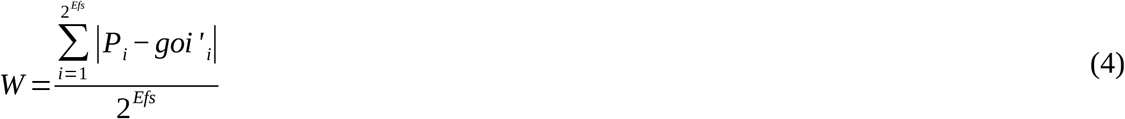

### Evolutionary protocol

In each generation, the alleles encoding for the GRN undergo point mutations: in a given generation, each element *B_ij_* changes its value to *B_ij_*+*v* with a probability *p*=*1/N_g_*, being *v*~*U*(−0.1,0.1), which ensures that mutational effects are independent of the GRN size. We do not mutate the *G* vector, which is initialized to the same initial value as in *t*=0 in each generation (we keep it constant because we are interested in the environmental-phenotype properties of the system, rather than in those properties derived from the more widely studied genotype-phenotype-maps *sensu* [13]). Each mutational event uniquely determines a mutant phenotype, whose probability of being fixed is proportional to the selective advantage of that mutation. Thus, we confront again all the clone sub-populations bearing the mutant allele with all possible environments, recording the mutant’s fitness *Wm*. If *Wm* ≥ *W*, then the new mutation is fixed.

We use this hill-climbing evolutionary algorithm because our aim is to provide a phenomenological model of the evolution of structured phenotypic plasticity, rather than quantifying realistic evolutionary rates or other important population or genetic parameters (e.g. magnitude of selection coefficients, mutation probabilities, population structure, epistatic effects) [20]. In doing so, we assume that the selection coefficient is sufficiently large that beneficial mutations get deterministically fixed and deleterious mutations do not. Although we acknowledge that more stochastic evolutionary scenarios may have qualitatively different dynamic properties, previous work suggest that both approaches yield consistent results [52].

### Experiment 2. Exploration of the morphospace of cell plasticity

In this experiment we performed an unbiased scanning of the parameter space to investigate how the different types of plasticity are spread over the theoretical space of all possible GRNs and how abundant each multi-dimensional reaction norm is. To do this, we generated a large number (≈10^5^) of random GRNs with a size of (3<*n_g_*<24), so that two of their genes were environmentally sensitive to *Ef_1_* and *Ef_2_* and a third, different gene was set as *goi*. In order to obtain representative networks for different values of the parameters considered (Average strength of connection and percentage of non-zero connections, which are proxies for the *L*_1_ and *L*_2_ regularisations parameters respectively, see costly connections below) the values of the *B_ij_* matrix for a given GRN were given, with *p*=*U*~(0,1), non-zero values randomly extracted from a normal distribution N~(U~(−1,1),σ) with σ=0.1. That way we avoid the convergence of the average absolute strength (*L*_1_) to a value of 0.5, derived from the central limit theorem.

### Experiment 3: Evolvability assays

We further assess if there exist some bias in the evolutionary transitions between the different forms of cell plasticity. Specifically, we focused on forms of plasticity associated with logical functions of different complexity Ω, that is, we checked if (*p*(*a* → *b*)=*p*(*b* → *a*) ∀ Ω_a_>Ω_b_). We first trained a number of GRNs (*n*=30) to produce a specific multi-dimensional reaction norm with an associated complexity of Ω_a_. When the GRN reaches a *W*=1, the simulation is stopped. Then we clone the trained network and force each clone to evolve again towards a new form of plasticity, now characterized by a complexity Ω_b_. We recorded the inverse of the number of generations required to attain again a *W*=1 as a proxy of the likelihood of the transition *a* → *b*, which can be interpreted as the system’s evolvability.

We checked if the results of these evolvability assays could be explained by the differences in the relative frequency of each type of cell plasticity in the GRN space (see Experiment 2). We calculated the expected long-term distribution of each type of plasticity derived from the probability transitions by iterating the differential equation (5), and compared the steady-state values in *t*_→∞_ with those resulting from the experiment 2.

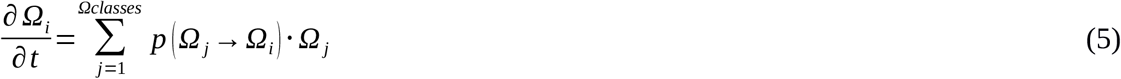

Where Ω_i_ is the relative frequency of the type of complexity *i* (amongst the Ω_classes_) and p(Ω_j_ → Ω_i_) is the transition probability between the Ω_j_ and the Ω_i_ classes.

### Experiment 4. Generalisation

In experiment 1, natural selection can find arbitrary forms of cell plasticity, but this required cells to be exposed to all possible combinations of environmental factors (complete information). In this experiment 4a, we instead expose plastic cells to a subset of all possible environments (incomplete information) and test under which circumstances they are able to generalize a suitable phenotypic response for the remaining ones. We refer to this subset of environments as the training set (*Ts*<*EFs*).

We evolve the system in a training phase until the maximum possible fitness under that specific *Ts* size (so that cells produce the required phenotypic states for each of the *Ts* environments). For each replicate, the composition of the training set is randomly chosen from the set of *2^EFs^* possible environments.

In the second (test) phase of the experiment, we expose the trained GRNs to the remaining *2^EFs^–Ts* environments, which are utterly novel to the cells. The fitness W is calculated now over all the *2^EFs^* environments. If the cells are able to generalize from their past evolutionary experience, the performance of the plastic cells in these new environments will be better than at random: *W*≥(2^*EFs*^−*Ts*)/2 (Figs 6A-B, and S1 Fig, green lines).

We further perform another generalisation experiment (Experiment 4b) using an explicit spatial 2D field of 1250 cells (25x50 sub-hexagonal grid) instead of a space-less implementation (Fig 7). In this case, the target developmental pattern was chosen amongst a large set of random patterns resulting from the integration of three morphogens (*EFs*) according to random thresholds. The pattern, resembling an arthropodian early embryo, was specifically chosen to be generated from simple logical functions and visually recognizable as a biological structure. The spatial distributions (*xy* coordinates) of the morphogens were drawn from simple and biologically realistic gradients: a Turing-like stripped pattern ([Ef1]_xy_=sen(x·k)), an isotropic radial gradient from a punctual signalling centre ([Ef2]_xy_=(x^2^+y^2^)^*1/2*^) and a lateral diffusion from the antero-posterior midline ([Ef3]_xy_=y^1/2^)

The combination of these morphogens generates the target developmental pattern according to the composite logical function AND(OR(AND(1,2),AND(2,3)),1) (Ω≪√(*EFs* ·2^*EFs+1*^)). This function is associated with a truth table of length *n*=32 (an arbitrary threshold was applied to the continuous morphogen gradients in order to reduce the length of the truth table). For the training phase, each individual cell was exposed to a random subset of this table (*TS*<32). For the test phase, every cell is exposed to the whole information (*TS*=32). We contrast these results with another scenario in which the missing information in the test phase is generated randomly (Fig 7B, S1 Fig).

### Experiment 5. Costly connections

In computer science, there are several procedures aimed to improve the performance of learning and generalisation experiments. Our aim in this experiment is to assess if they are also applicable in the context of cell plasticity. Amongst these procedures, a widespread one consist basically in limiting the complexity that the network can attain. It can be easily done by making the GRN connections costly, with a direct effect on fitness:

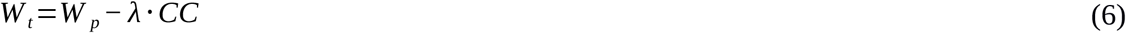

Where *Wp* is the partial fitness scored as in Equation (4) and λ is the relative weight of the cost of connections (*CC*) in determining the total fitness *Wt*. In turn, *CC* can be implemented in two ways, called respectively *L*_1_ and *L*_2_ regularisation. In the *L*_1_ regularisation, *sparse* connectivity is favoured by applying a direct selective pressure in the number gene-gene interactions, which decreases the number of connections in the GRN [26]. Li is implemented as the sum of absolute magnitudes of all regulatory interactions:

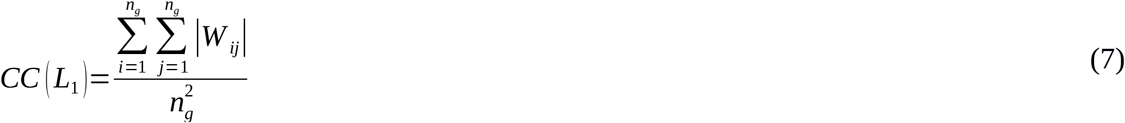

In the *L*_2_ regularisation, *weak* connectivity is favoured by applying a direct selective pressure in the strength of gene-gene interactions, which results in regulatory interactions of small magnitude [26]. *L*_2_ is implemented as a the sum of the squares of the magnitudes of all regulatory interactions:

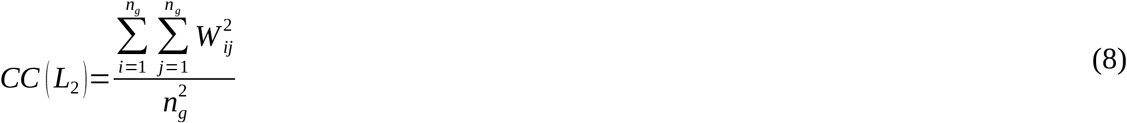

## Acknowledgements

The authors thank T. Uller, C. Thies, A. Rago, D. Prosser, J. Caldwell, J.F. Nash, L.E. Mears and I. Hernando-Herraez for helpful discussion and comments.

